# Soil aggregates reveal tree species and land-use legacy effects on early carbon storage pathways during reforestation

**DOI:** 10.64898/2026.09.22.753307

**Authors:** Courtney E. Mathers, Hilary Rose Dawson, Emily Huckstead, Lucas C. R. Silva

**Affiliations:** Environmental Studies Program, University of Oregon, Eugene, Oregon, USA; Department of Biological Sciences, University of Bergen, Bergen, Norway; Institute of Ecology and Evolution, University of Oregon, Eugene, Oregon, USA; Department of Biology, University of Oregon, Eugene, Oregon, USA

**Keywords:** reforestation, soil carbon, soil aggregates, aggregate stability, land-use legacy, mycorrhizal strategy, restoration monitoring

## Abstract

Reforestation is a leading natural climate solution, but bulk soil organic carbon (C) often responds slowly, obscuring early belowground change. We tested whether soil aggregates reveal early structural reorganization that affects C retention pathways at a three-year-old experimental reforestation planting established on former pasture in Oregon, USA. We sampled soils at 0-20 and 20-40 cm beneath incense cedar (*Calocedrus decurrens*; arbuscular mycorrhizal [AM]), black cottonwood (*Populus trichocarpa*; AM and ectomycorrhizal [EcM]), ponderosa pine (*Pinus ponderosa*; EcM), and treeless controls. We measured bulk soil C concentration and C:N, aggregate size distribution and mean weight diameter (MWD), fraction-associated C, and, in a subset of surface aggregate fractions, natural-abundance δ¹³C to evaluate soil C pools, physical structure, C distribution, and C processing. After three years, reforestation did not produce significant differences in bulk soil C between planted trees and treeless controls. In surface soil, MWD averaged 36% higher under incense cedar and 58% higher under black cottonwood than under controls, whereas ponderosa pine remained similar to controls. The clearest treatment differences in C distribution occurred in macroaggregates. Incense cedar and black cottonwood had higher large (>2000 µm) macroaggregate-associated C than controls, while black cottonwood combined a greater proportion of large macroaggregates with lower C concentrations, indicating that structural development and C accumulation were partly decoupled. Species patterns were broadly consistent with stronger early aggregate responses under AM-compatible species in former-pasture legacy conditions; by contrast, the EcM-associated ponderosa pine remained closer to treeless controls across multiple aggregate measures. Treatment effects weakened with depth and were limited in microaggregates (250-53 µm) and silt-and-clay (<53 µm) fractions. Smaller aggregate fractions were progressively enriched in δ¹³C, consistent with greater C processing in protected fractions, but treatment-level isotope differences had not yet emerged. Soil aggregates showed species-associated belowground reorganization, revealing how tree identity and former-pasture legacy may shape early soil C organization before consistent differences emerged in bulk soil C. Aggregate measures can complement bulk soil C measurements in restoration monitoring as early indicators of belowground soil C trajectories shaped by tree species identity and land-use legacy.

**Open Research Statement:** All raw data and metadata are provided for peer-review with Dryad at xxxxx, where it will be permanently archived for public access upon acceptance.

## Introduction

Forests are a major component of the global carbon (C) cycle, and reforestation has been identified as a leading natural climate solution for climate change mitigation through increased C storage (Griscom et al., 2017). Although reforestation produces substantial gains in aboveground biomass relatively quickly, belowground C responses are slower to emerge and more variable (Li et al., 2012; Paul et al., 2002; Paul et al., 2016). This variability reflects the sensitivity of soil C trajectories to climate, local soil conditions, previous land use, and tree species identity (Laganière et al., 2010; Mayer et al., 2020; O’Kelley et al., 2025; Paul et al., 2002). As a result, young plantings may show little net change in soil C even while developing trees alter organic matter (OM) inputs, microbial activity, and soil physical structure. Reliance on bulk soil C alone may miss early belowground treatment differences relevant to restoration monitoring. More responsive indicators are needed to evaluate early belowground change and identify which species or site conditions are beginning to shape soil C trajectories (Gatica-Saavedra et al., 2023).

Soil aggregates provide a responsive indicator of early soil C trajectories because their formation and turnover respond rapidly to roots, microbial activity, and soil disturbance (Rillig et al., 2017; Totsche et al., 2018). Aggregates are structural units that form as mineral particles, OM, roots, fungal hyphae, and microbial products bind together (Tisdall & Oades, 1982). Most topsoil C is associated with aggregates (Jastrow, 1996), and aggregate structure plays a central role in organizing OM into physical habitats that regulate microbial access to C and physically protect OM from decomposition (Jastrow et al., 2007; Six et al., 2004; Wilpiszeski et al., 2019). Aggregate formation and breakdown also influence water movement and erosion resistance (Barthès & Roose, 2002), making aggregate size distribution and stability common indicators of soil physical condition during vegetation recovery or land management change (Bronick & Lal, 2005; Bagnall et al., 2023; Hamza et al., 2025). In young reforestation settings, these aggregate-based measures may reveal developing soil structure and C storage pathways before bulk soil C pools show clear change, but their value as early indicators remains underexplored (Gatica-Saavedra et al., 2023).

Different aggregate-based measurements provide complementary information about soil C distribution and processing. Macroaggregates (>250 µm) are relatively dynamic sites of recent C incorporation and turnover, whereas microaggregates (<250 µm) and mineral-associated fractions are associated with longer-term C protection (Six et al., 2000; Six et al., 2004; Tisdall & Oades, 1982). Aggregate stability and size distribution indicate changes in soil physical structure and the balance between aggregate formation and disruption, while aggregate-associated C and fraction-specific C concentrations show where C is distributed among structural fractions (Rieke et al., 2022; Six et al., 2004). Natural-abundance δ^13^C adds information about C processing within these fractions because microbial transformation preferentially removes ^12^C through respiration, leaving remaining OM relatively enriched in ^13^C (Gunina & Kuzyakov, 2014; Werth & Kuzyakov, 2010). Smaller, more protected aggregate fractions often show higher δ^13^C values than larger aggregates, consistent with greater microbial processing and relative stabilization of C in these fractions (Liu et al., 2018; Wang et al., 2025). Together, these measurements provide integrated indicators of developing soil structure, C distribution, and potential pathways of C stabilization, rather than direct substitutes for soil C stock measurements.

In reforestation contexts, tree species identity may shape early aggregate responses because roots, litter inputs, growth strategy, and mycorrhizal fungi influence the quantity and form of OM entering soil. Roots supply organic inputs and physically enmesh soil particles, while mycorrhizal fungi are especially important for aggregate stabilization because hyphae extend beyond root surfaces and bind particles into water-stable structures (Miller & Jastrow, 1990; Rillig & Mummey, 2006; Rillig et al., 2015). Mycorrhizal strategy may therefore provide one mechanism by which tree species differ in their early effects on aggregate-mediated soil C dynamics (De Geode et al., 2025; Lehmann et al., 2020). Arbuscular mycorrhizal (AM) fungi grow into root tissues and are common among grasses and herbaceous plants, whereas ectomycorrhizal (EcM) fungi form sheaths around fine roots and are common among many temperate forest trees (Brundrett & Tedersoo, 2018). AM and EcM fungi differ in hyphal traits, nutrient acquisition strategies, and interactions with surrounding microbial communities, which may lead to different effects on aggregation and soil C dynamics (Averill et al., 2014; Qiang et al., 2023; Shah et al., 2016). Experimental reforestation sites provide an opportunity to test whether tree species with contrasting mycorrhizal associations diverge in aggregate-mediated soil C dynamics under common field conditions.

Former pasture provides a useful context for testing mycorrhizal legacy effects. Natural succession from grassland or shrubland toward forest is often accompanied by shifts in mycorrhizal dominance, with AM plants common in earlier herbaceous stages and EcM trees becoming more prominent in later forest stages (Qiang et al., 2023). Reforestation on former pasture compresses this belowground transition by introducing tree seedlings into soils still shaped by herbaceous vegetation and AM-associated microbial communities. This legacy may be especially important for EcM tree species, whose establishment can be limited when compatible fungal communities are absent or reduced (Nuñez et al., 2009; Wang et al., 2023). These legacy effects may influence tree survival, root and fungal development, and aggregate-mediated soil C dynamics during early reforestation.

We used a three-year-old experimental reforestation planting established on former pasture to test whether aggregate-based measurements detect early belowground differences that are not apparent in bulk soil C. We compared soils beneath three planted tree species with contrasting reported mycorrhizal associations and adjacent treeless controls, while recognizing that species identity and mycorrhizal effects could not be separated experimentally. Because early belowground effects depend on successful tree establishment, we also evaluated survivorship across the broader site and tree height among the surviving individuals sampled for soil analyses. We hypothesized that treatment differences would emerge first in surface-soil macroaggregate distribution, stability, and associated C, before becoming detectable in bulk soil C. We further expected δ¹³C enrichment toward smaller fractions, consistent with progressively processed OM within the aggregate hierarchy. Given the influence of mycorrhizal fungi on tree growth and soil structure, we expected AM trees to show stronger early aggregate responses than EcM trees under the site’s herbaceous AM-legacy conditions. By integrating measures of aggregate stability, aggregate size distribution, aggregate-associated C, and δ^13^C, this study tests whether soil aggregates can reveal early reforestation effects on C dynamics while clarifying how tree species identity and land-use legacy shape initial belowground carbon pathways.

## Methods

### Study site and sampling design

The study was conducted at an experimental riparian reforestation site along the McKenzie River in Oregon’s Willamette Valley (44.063043, −122.931603). The region has a Mediterranean climate, with warm dry summers and cool wet winters, has a mean annual temperature of 11.5 °C, and receives a mean annual precipitation of 1145 mm (PRISM Group, Oregon State University, 2026). The site features Mollisols with an average soil pH of 6.62 ± 0.81 (SD). Riparian hardwood forest characterized the site in the 1880’s according to local land survey models (Christy & Alverson, 2011). The site was managed as agricultural pasture from at least the 1930s until the early 1960s and had been left fallow for approximately 60 years prior to reforestation. In February 2021, planting furrows were established and tree seedlings were planted at approximately 12-ft spacing. No mycorrhizal inoculum was added at the time of planting. Tree species included in this study were EcM conifer ponderosa pine (*Pinus ponderosa*)(Walker et al., 2010); AM conifer incense cedar (*Calocedrus decurrens*) (Kough et al., 1985); and black cottonwood (*Populus trichocarpa*), a broadleaf species reported to host both AM and EcM fungi (Nash et al., 2025). These associations were not experimentally manipulated or measured at the site; consequently, tree species was treated as the experimental factor, and mycorrhizal strategy was considered only as an explanation for species-specific patterns. Although the reforestation site included additional planted species and extended beyond the area sampled here, soil sampling for this study was restricted to a topographically uniform subsection (4.8 ha) of the planting. This design minimized landscape-scale variation in soil conditions, standardized when the trees were planted, and allowed differences among the focal tree treatments and treeless controls to be evaluated within a common soil template.

We assessed sapling survivorship in July 2021 and again in April of 2023 using a three-category field score reflecting likelihood of survival to the following year (3 = strong chance of surviving to next year, 2 = moderate chance of surviving to next year, 1 = appears deceased). For the three focal tree species, we summarized survivorship from the 2021 survey to the 2023 survey as the proportion of individuals recorded as alive in 2023. Individuals with 2023 survival scores of 2 or 3 were classified as alive, and individuals with scores of 1 were classified as dead. Survivorship was summarized across the broader planting to provide context for species-level differences in establishment (Table 1).

**Table 1:** Focal tree species differed in mycorrhizal association, survivorship across the broader planting, and height of healthy surviving individuals randomly sampled for soil analyses.

| Tree Species | Mycorrhizal association | Planted trees (n) | Survivorship (%) | Sampled trees (n) | Sampled tree height (cm), mean ± SD |
| --- | --- | --- | --- | --- | --- |
| Incense cedar | AM | 433 | 43.4 | 15 | 93.5 ± 21.2 |
| Black cottonwood | AM/EcM | 573 | 26.4 | 15 | 137.3 ± 79.1 |
| Ponderosa pine | EcM | 244 | 12.3 | 15 | 74.0 ± 19.1 |

**Notes:** Survivorship is reported as the percentage of focal individuals alive in 2023, six months prior to soil sampling. Tree height values are means ± SD.
**Abbreviations:** AM, arbuscular mycorrhizal; EcM, ectomycorrhizal; SD, standard deviation.

In October 2023, we collected soil samples (n = 116) with a soil core sampler positioned 20 cm from the base of each sampled sapling to target the near-root zone while avoiding direct disturbance of the stem base. Separate samples were collected from 0-20 cm and 20-40 cm depths to assess whether treatment effects differed between surface and deeper soil layers. Fifteen healthy individuals of each tree species were randomly selected for sampling, and 13 control samples were collected at both depths from adjacent treeless locations within the same field. Only healthy surviving trees with survivorship scores of 3 in the 2023 survey were sampled to minimize confounding effects of poor establishment. The height of each sampled tree was also recorded in cm (summarized in Table 1). All tree and control samples were collected from within furrows because furrow establishment likely influenced soil structure. Fresh soils were passed through an 8-mm sieve prior to air-drying and were then stored at room temperature until analysis.

### Aggregate fractionation and aggregate stability

We separated each bulk soil sample into four aggregate-size fractions: large macroaggregates (> 2000 µm), small macroaggregates (2000–250 µm), microaggregates (250–53 µm), and silt-and-clay particles (< 53 µm). We conducted fractionation following a modified Elliott (1986) procedure with a Royal Eijkelkamp wet sieving apparatus (Giesbeek, The Netherlands). Briefly, a 4-g subsample of air-dried soil was placed on the 2000-µm sieve and saturated with deionized water for 3 min to induce slaking. After slaking, the sieves were oscillated vertically in water 100 times over 3 min. Material retained on each sieve was collected, and the remaining suspension was transferred sequentially to the next smaller sieve. This process was repeated through the 250-µm and 53-µm sieve classes. Fraction samples were oven-dried at 60 °C and weighed. Rock fragments (> 2000 µm) were removed from the large macroaggregate fraction, and their mass was subtracted from both the recorded large macroaggregate mass and total dry soil mass used in subsequent calculations.

We calculated aggregate mass proportions as the dry mass of each fraction divided by the total dry soil mass used in wet sieving. We quantified aggregate stability as mean weight diameter (MWD), a widely used index of aggregate stability in which larger values indicate a greater proportion of soil in larger water-stable aggregates (Reike et al., 2022). MWD was calculated as:

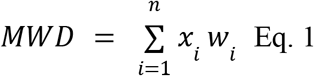

where *x_i_* is the mean diameter of the sieve size class retained in fraction *i* and *w_i_* is the ratio of stable aggregate mass to total dry soil mass for that fraction.

### Elemental and isotopic analyses

We analyzed all bulk soil samples (n = 116; n = 58 per depth) and aggregate fractions (n = 464; n = 58 per size fraction × depth combination) for total C and nitrogen (N) concentrations using a FlashSmart Elemental Analyzer (Thermo Scientific, Grand Island, NY). All bulk samples were first sieved to 2000 µm and ground to a fine powder before combustion analysis. Preliminary site analyses indicated no inorganic soil carbonates, so total C represents total soil organic C (SOC) throughout. Bulk soil C concentration was converted from percent C to g C kg^−1^ soil to express C mass per unit dry soil mass. Bulk soil C:N was calculated from bulk soil C and N concentrations, and aggregate-fraction C:N was calculated from fraction-specific C and N concentrations. For aggregate size fractions, we distinguished between fraction-specific C concentration and fraction-associated C amount. Fraction-specific C concentration refers to the C concentration of a given aggregate size fraction. Fraction-associated C (g C kg^−1^ bulk soil) refers to the amount of C contributed by that size fraction per unit mass of bulk soil and was calculated by multiplying the fraction’s mass proportion by its fraction-specific C concentration (Zhao et al., 2023). Estimated total aggregate-associated C was calculated by summing the fraction-associated C amounts across all four aggregate fractions.

Stable C isotopes (^13^C/^12^C) were measured for a subset of 0-20 cm samples across the four aggregate fractions (n = 64; n = 4 per treatment × size fraction combination). This subset included four sets of aggregate fractions that were randomly selected within each treatment. Aggregate-fraction C:N was evaluated alongside δ¹³C as an additional indicator of C processing across aggregate fractions. C isotope ratios were measured at the Stable Isotope Facility, University of California, Davis using an Elementar vario EL cube elemental analyzer interfaced to an Elementar VisION isotope ratio mass spectrometer. Isotope ratios are expressed as δ^13^C relative to Vienna Pee Dee Belemnite (V-PDB), and the standard deviation for replicate reference materials was ± 0.12‰. We evaluated δ13C, together with aggregate-fraction C:N, as indicators of C processing and potential C stabilization across aggregate fractions (Scartazza et al., 2023).

### Statistical analyses

All analyses were conducted in R (v4.5.2; R Core Team, 2025). Data processing and visualization were conducted using tidyverse packages (Wickham et al., 2019). Treatments were defined as the four vegetation categories sampled at the site: treeless control, incense cedar, black cottonwood, and ponderosa pine. Base R functions were used for ANOVA, Welch’s ANOVA, chi-square tests, linear models, Pearson correlations, and associated pairwise comparisons. We evaluated variable distributions and model residuals prior to analysis and applied log or logit transformations where needed to improve residual normality and homogeneity of variance. Aggregate proportion variables were analyzed on the logit scale, although raw values are presented in figures for interpretability. MWD was analyzed and presented on the log scale. Statistical significance was assessed at α = 0.05 throughout.

Survivorship among the three focal tree species was evaluated using a chi-square test of independence. Sampled tree height was compared among species using Welch’s ANOVA on log-transformed height, followed by pairwise Welch t-tests with Benjamini-Hochberg correction. Response variables measured once per bulk soil sample, including bulk soil C concentration and bulk soil C:N, aggregate stability, aggregate mass proportions, and total aggregate-associated C, were analyzed using ANOVA. Aggregate proportions and aggregate-associated C and C:N were analyzed separately for each aggregate fraction and depth, with depth-specific treatment comparisons evaluated using Tukey’s HSD tests. Variables compared across multiple aggregate fractions from the same bulk soil sample, including fraction-specific C and N concentrations, aggregate-fraction C:N, and δ13C, were analyzed using linear mixed-effects models with sample identity as a random intercept using the lme4 package (Bates et al., 2015). Fixed effects for mixed models were evaluated using the lmerTest package (Kuznetsova et al., 2017), and Tukey-adjusted pairwise comparisons from mixed-effects models were calculated using the emmeans package (Lenth, 2026). Relationships among aggregate stability (MWD), bulk soil C, bulk soil C:N, and tree height were evaluated using Pearson correlations and linear models that included depth and treatment. Interaction models were used to test whether these relationships varied by depth, treatment, or their interaction.

## Results

### Tree establishment differed among focal species

Survivorship differed strongly among the three focal tree species across the broader restoration planting (χ² = 77.76, df = 2, *p* < 0.001; Table 1). Incense cedar had the highest survivorship (43.4%), followed by black cottonwood (26.4%) and ponderosa pine (12.3%). Among surviving trees sampled for soil analyses, height also differed among species based on Welch’s ANOVA of log-transformed height (*F*_2,54.09_ = 11.34, *p* < 0.001). Black cottonwood was tallest, followed by incense cedar and ponderosa pine, with all pairwise comparisons significant (all *p* ≤ 0.039; Table 1). These species-level differences in establishment and early growth provide context for the aggregate and soil C patterns reported below.

### Bulk soil C did not differ between three year old planted trees and treeless controls

Reforestation did not produce significant differences in bulk soil C concentration between planted trees and treeless controls at either soil depth, although differences among planted species were apparent (Figure 1A). Bulk soil C was greater in the 0-20 cm layer than in the 20-40 cm layer across all treatments (*F*_1,107_ = 39.71, *p* < 0.001), with mean values (± SD) of 25.1 ± 5.6 g C kg^−1^ soil and 19.3 ± 4.8 g C kg^−1^ soil, respectively. In the 0-20 cm layer, bulk soil C differed overall among treatments (*F*_3,54_ = 2.93, *p* = 0.042), but no pairwise contrasts were significant in Tukey’s HSD tests. Surface soil patterns nevertheless suggested lower bulk soil C in black cottonwood soils than in incense cedar (*p* = 0.053) and control soils (*p* = 0.076). In the 20-40 cm layer, treatment differences were more pronounced (*F*_3,53_ = 3.51, *p* = 0.021), with incense cedar soils containing greater bulk soil C than black cottonwoods (*p* = 0.013), while control and ponderosa pine soils were intermediate and did not differ significantly from either incense cedar or black cottonwood (all *p* ≥ 0.162; Figure 1A).

**Figure 1:**
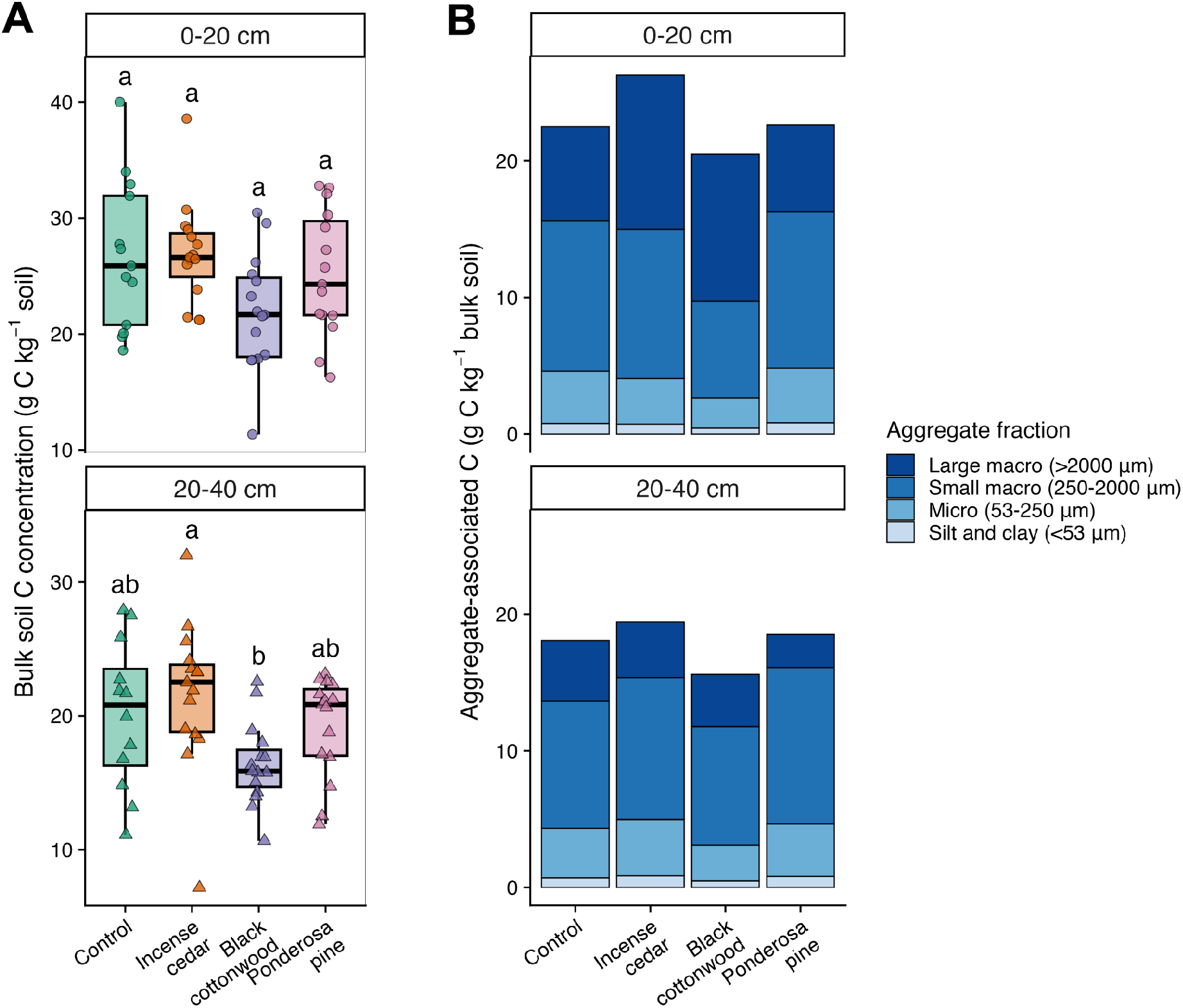
Bulk soil C concentration and mean aggregate-associated C under control, incense cedar, black cottonwood, and ponderosa pine soils at 0-20 cm and 20-40 cm depths. Mycorrhizal associations are incense cedar = AM, black cottonwood = AM/EcM, and ponderosa pine = EcM. (A) Bulk soil C concentration (g C kg^−1^ soil) shown as boxplots with individual sample values; different letters indicate significant differences among species within each depth based on pairwise Tukey’s HSD tests. (B) Mean aggregate-associated C (g C kg^−1^ bulk soil); bar totals represent the estimated bulk soil C pool calculated by summing mean fraction-associated C amounts, with colors indicating the contribution of each aggregate size fraction.

### Tree species differed in aggregate-associated C amounts and concentrations

Aggregate-associated C showed clearer treatment differences than bulk soil C, especially in larger surface soil fractions. Summed fraction-associated C recovered approximately 93% of the soil C measured independently in bulk soil, indicating generally strong analytical recovery through the fractionation procedure. Aggregate-associated C was concentrated primarily in the large and small macroaggregate fractions, whereas microaggregates and the silt-and-clay fraction contributed comparatively little to total C mass (Figure 1B). In the 0-20 cm layer, large macroaggregate C differed among treatments in the surface soil (*F*_3,54_ = 4.09, *p* = 0.011), with mean values under incense cedar and black cottonwood 64% and 57% greater than the treeless control, respectively, although pairwise differences were less clear. Small macroaggregate C also differed among treatments (*F*_3,54_ = 6.69, *p* < 0.001), but followed a different pattern: black cottonwood soils had the lowest small macroaggregate-associated C, 36% below controls, and were significantly lower than control, incense cedar, and ponderosa pine soils (all *p* < 0.01; Appendix S1: Table S1). Treatment differences in larger aggregate-associated C pools were weaker in the 20-40 cm layer, where large macroaggregate C did not differ among treatments and small macroaggregate C showed only a marginal treatment effect (*F*_3,54_ = 2.32, *p* = 0.086; Appendix S1: Table S1).

Fraction-specific C concentrations (%) provided additional context for these aggregate-associated C patterns. Across treatments, large macroaggregates were the most C-rich fraction, even though small macroaggregates often held the largest share of total C mass (Appendix S1: Table S2). Fraction-specific C concentration differed among aggregate fractions overall (*F*_3,324_ = 94.40, *p* < 0.001), and the effect of fraction varied by depth (*F*_3,324_ = 5.29, *p* = 0.001). At both depths, large macroaggregates had the highest mean C concentration, averaging, 2.70 ± 0.47% C in the 0-20 cm layer and 2.42 ± 0.57% C in the 20-40 cm layer (mean ± SD). Large macroaggregate C concentration was significantly greater than all other fractions at both depths (all p < 0.001). Fraction-specific C concentration also differed among treatments overall (*F*_3,108_ = 13.07, *p* < 0.001). Black cottonwood had the lowest mean C concentration in each aggregate fraction (Appendix S1: Table S2) and was lower than control (*p* = 0.009), incense cedar (*p* < 0.001), and ponderosa pine (*p* < 0.001). Because fraction-associated C was calculated from both aggregate mass and fraction-specific C concentration, the black cottonwood pattern reflected both its greater large macroaggregate proportion and its lower fraction-specific C concentrations. Incense cedar had the highest mean fraction-specific C concentration, although it was only marginally higher than the control (*p* = 0.051) and did not differ significantly from ponderosa pine (*p* = 0.381).

### Surface aggregate stability varied among treatments

Aggregate stability, quantified as log-transformed mean weight diameter (MWD), differed among treatments in the 0-20 cm layer (*F*_3,54_ = 5.43, *p* = 0.002; Figure 2A). Black cottonwood had the strongest surface-soil aggregate stability response despite having lower bulk soil C than incense cedar. Relative to treeless controls, average MWD was approximately 36% higher under incense cedar and 58% higher under black cottonwood, whereas ponderosa pines remained similar to control stability values. Black cottonwood soils had 58% higher MWD than control (*p* = 0.03) and 68% higher MWD than ponderosa pine soils (*p* = 0.006), while incense cedar was intermediate and did not significantly differ from control (*p* = 0.177), black cottonwood (*p* = 0.842), or ponderosa pine (*p* = 0.052; Figure 2A). Treatment effects on MWD were confined to the 0-20 cm layer, as MWD did not differ among treatments in the 20-40 cm layer (*F_3,54_* = 1.16, *p* = 0.335).

**Figure 2:**
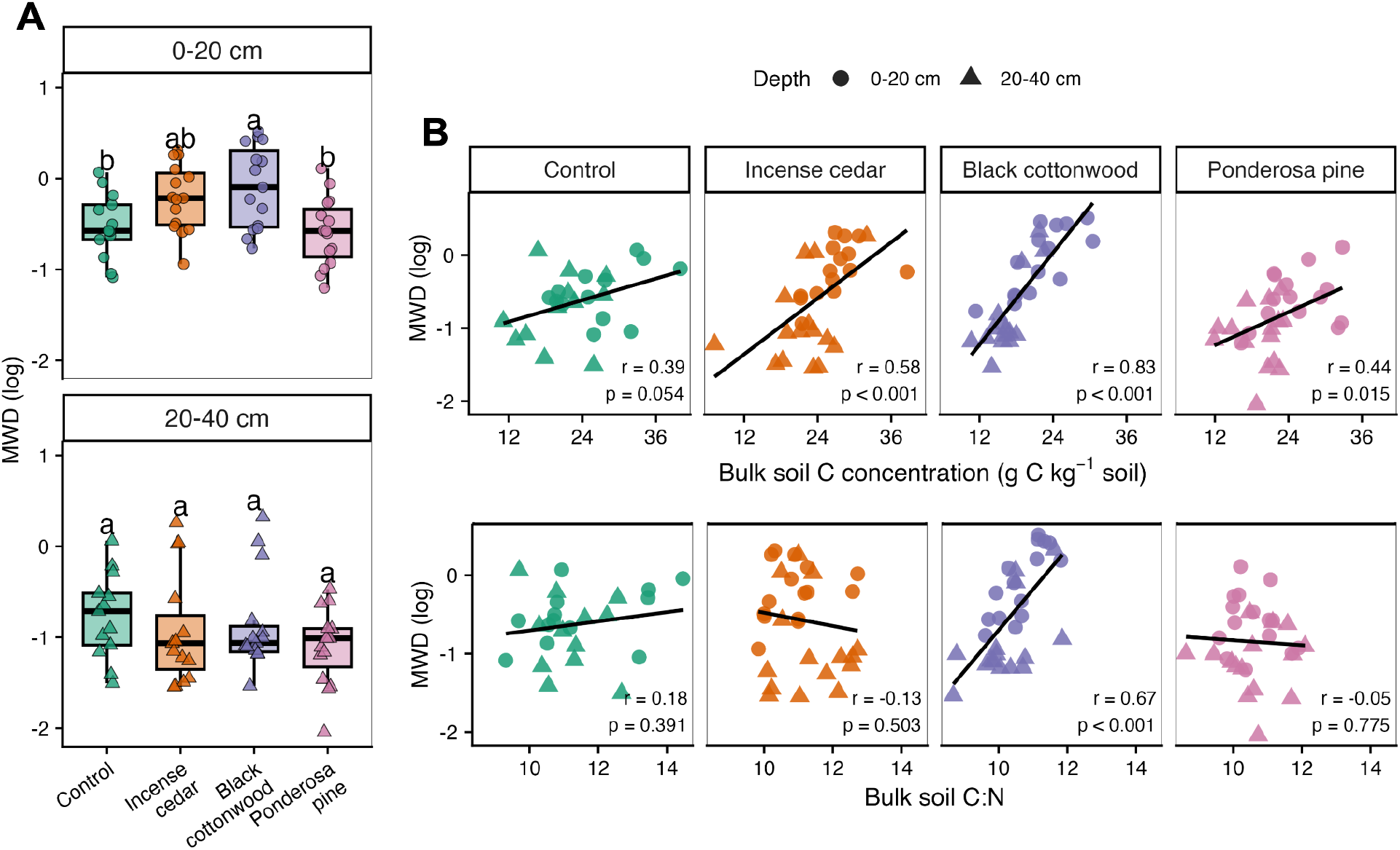
Aggregate stability and its relationships with bulk soil C and bulk soil C:N under control, incense cedar, black cottonwood, and ponderosa pine soils. (A) Aggregate stability (log transformed MWD) across treatments and depths. Boxplots show medians, interquartile ranges, and individual sample values; different letters indicate significant differences among treatments within each depth based on pairwise Tukey’s HSD tests. (B) Relationships between log transformed MWD and bulk soil C concentration (g C kg^−1^ soil) and bulk soil C:N. Points are colored by treatment and shaped by depth; black lines show treatment-specific linear fits. Pearson correlation coefficients (*r*) and associated *p* values are shown in each panel.

Treatment differences in surface-soil aggregate stability were expressed primarily through shifts in the balance between large and small macroaggregate pools, with incense cedar and black cottonwood showing stronger shifts toward larger aggregates than ponderosa pine or treeless controls. Because MWD reflects the distribution of soil among aggregate-size classes, we examined the fraction proportions underlying stability patterns (Appendix S1: Figure S1). In the 0-20 cm layer, fraction proportions differed among treatments, with the clearest differences occurring in the macroaggregate fractions. Large and small macroaggregate proportions both differed among treatments (*F_3,54_* = 5.66, *p* = 0.002, and *F_3,54_* = 3.82, *p* = 0.015, respectively). Based on treatment means, the proportion of bulk soil in large macroaggregates was 48% higher under incense cedar and 79% higher under black cottonwood relative to treeless controls, whereas ponderosa pine remained similar to control values (Appendix S1: Figure S1). Large macroaggregate proportion was greater in black cottonwood than in control (*p* = 0.02) and ponderosa pine soils (*p* = 0.003), while small macroaggregate proportion was greater in ponderosa pine than in black cottonwood soils (*p* = 0.012). None of the fraction proportions differed significantly among treatments in the 20-40 cm layer.

Variation in aggregate stability was more closely related to bulk soil C concentration than to bulk soil C:N, but the strength of these associations differed among treatments (Figure 2B). MWD was positively associated with bulk soil C concentration across samples (*r* = 0.48, *p* < 0.001), indicating that soils with greater physical stability also tended to contain more C (Appendix S1: Figure S2). This relationship was not explained solely by depth or treatment differences (*F*_1,109_ = 19.37, *p* < 0.001). However, the relationship strength varied among treatments (*F*_3,106_ = 5.57, *p* = 0.001), with the strongest positive relationships occurring in black cottonwood and incense cedar soils (Figure 2B). By contrast, MWD was not clearly related to bulk soil C:N (*r* = 0.15, *p* = 0.104), although the MWD-C:N relationship also varied among treatments (*F*_3,106_ = 4.55, *p* = 0.005) and was driven by a positive relationship in black cottonwood soils. All bulk soil C, N and C:N measurements are summarized in Appendix S1: Table S4. Sampled tree height was not significantly associated with MWD overall (*r* = 0.18, *p* = 0.087), but the height-MWD relationship varied among tree species (*F*_2,83_ = 5.72, *p* = 0.005), with a positive relationship under incense cedar only (Appendix S1: Figure S3).

### Smaller aggregate fractions were enriched in δ^13^C

Across treatments, δ^13^C values ranged from −28.25 to −26.62 ‰ and became progressively more enriched in the smaller aggregate fractions (Figure 3). Carbon isotope composition differed strongly among aggregate fractions (*F*_3,45_ = 20.86, *p* < 0.001), but not among treatments (*F*_3,12_ = 0.59, *p* = 0.631). The treatment × fraction interaction was also not significant (*F*_9,36_ = 0.69, *p* = 0.714), indicating that fraction-level δ13C patterns were broadly similar across treatments. Relative to large macroaggregates, small macroaggregates were enriched by 0.19‰, microaggregates by 0.36‰, and silt-and-clay fractions by 0.52‰. Pairwise comparisons showed that large macroaggregates were significantly more depleted than small macroaggregates (*p* = 0.039), microaggregates (*p* < 0.001), and silt-and-clay fractions (*p* < 0.001). Silt-and-clay fractions were also more enriched than small macroaggregates (*p* < 0.001), but did not differ significantly from microaggregates (*p* = 0.108). Aggregate-fraction C:N was evaluated using the full aggregate-fraction dataset, rather than only the isotope subset. Similar to the δ¹³C pattern, C:N generally decreased from macroaggregates to smaller fractions (Appendix S1: Table S3). Aggregate-fraction C:N differed among fractions overall (*F*_3,321_ = 9.60, *p* < 0.001), but not among treatments (*F*_3,108_ = 2.52, *p* = 0.062), with no treatment × fraction interaction (*F*_9,321_ = 1.27, *p* = 0.250; Appendix S1: Table S3).

**Figure 3:**
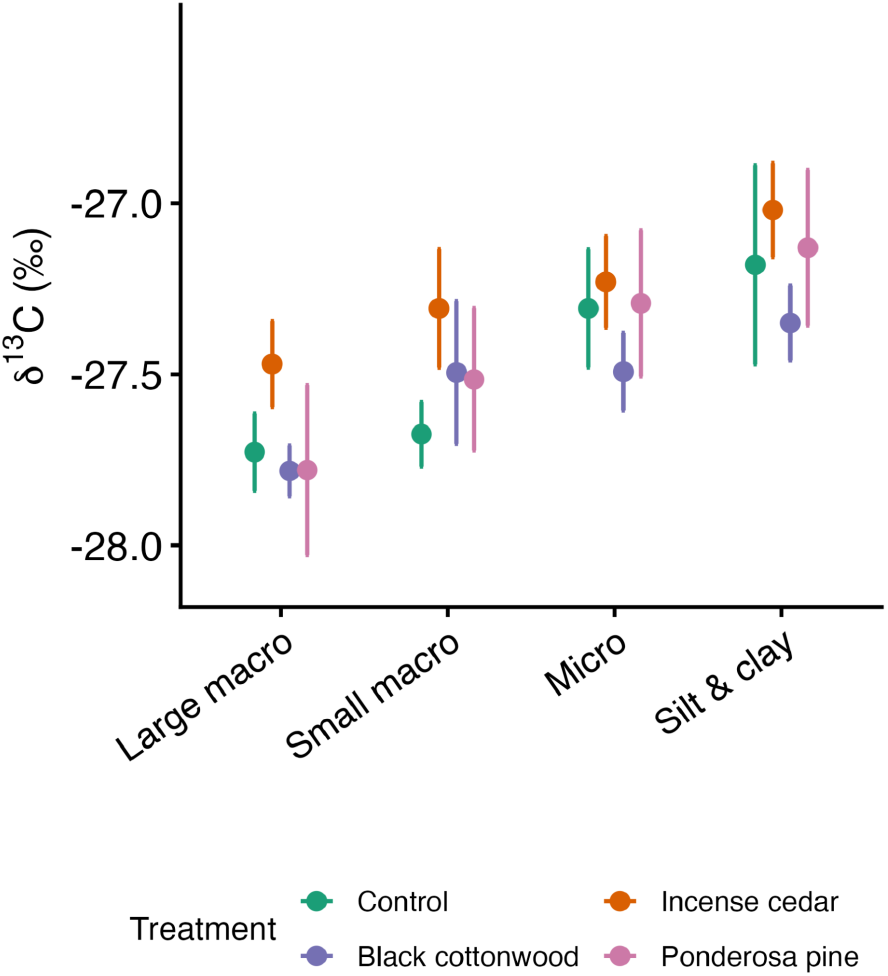
Carbon isotope composition (δ^13^C ‰) across aggregate fractions in the 0-20 cm soil layer under control, incense cedar, black cottonwood, and ponderosa pine. Points show treatment means and error bars show standard errors (total n = 64; n = 4 per species × fraction combination). Aggregate fractions are large macroaggregates (> 2000 µm), small macroaggregates (2000–250 µm), microaggregates (250–53 µm), and silt-and-clay particles (< 53 µm). Across treatments, δ^13^C values became progressively more enriched in the smaller aggregate fractions.

## Discussion

Bulk soil C responses to reforestation emerge slowly and unevenly, even though young trees are already altering belowground inputs, microbial activity, and soil structure. At a three-year-old riparian reforestation site established on former pasture, we tested whether soil aggregates provided evidence of early belowground change not captured by bulk soil C under shared land-use legacy conditions, and whether aggregate responses to reforestation varied among planted tree species with contrasting mycorrhizal associations. After three years, reforestation did not produce significant bulk soil C concentration differences relative to treeless controls, but tree treatments differed in aggregate-associated C, surface-soil aggregate stability, and the distribution of C among aggregate fractions. These differences were most pronounced in surface soil and in the larger aggregate fractions, indicating that reforestation altered the physical structure and distribution of soil C before producing detectable increases in bulk soil C relative to treeless controls. Responses also varied among tree species, providing partial support for our expectation that AM trees would show stronger early aggregate responses than EcM trees under former pasture legacy conditions. AM trees also had higher survivorship than EcM trees. These patterns were consistent with the influence of pasture legacy and species-specific reorganization of soil structure and belowground C retention pathways. Thus, the absence of bulk soil C gains did not indicate an absence of belowground change, but rather that early effects were expressed first as shifts in aggregate structure and the distribution of soil C among aggregate fractions.

These findings support our expectation that aggregates respond quickly to vegetation changes and reveal early reforestation effects not apparent from bulk soil C alone, providing new evidence that aggregate-based measures can help monitor belowground change in young restoration sites. The progressive enrichment of δ¹³C from macroaggregates toward microaggregates and silt-and-clay fractions added mechanistic context, indicating increasing microbial processing and relative protection of carbon across the aggregate hierarchy. Together, these patterns suggest that aggregate-scale carbon processing and structural reorganization can precede detectable gains in bulk soil C, making aggregates useful early indicators of developing C-retention pathways during reforestation.

### Aggregate responses precede bulk soil C gains

The absence of higher bulk soil C concentrations beneath planted trees after three years fits the expected timing of soil C development after reforestation. In western Oregon riparian systems, soil C has projected gains of 0.82 Mg C ha^−1^ yr^−1^ over the first 25 years after planting, yet soil C stocks are estimated to require more than a decade to significantly exceed baseline levels (O’Kelley et al. 2025). Soil C accumulation in the region is shaped by precipitation, edaphic conditions, and prior land-use history, reinforcing the need for site-specific monitoring (O’Kelley et al. 2025). Global-scale syntheses similarly show that soil C responses to afforestation and reforestation depend strongly on previous land use, tree species, soil properties, and stand age (Laganière et al., 2010; Paul et al., 2002). In this context, the concurrent differences in surface aggregate-associated C, MWD, and fraction-level δ¹³C patterns show that early belowground change was detectable in aggregate-based measures before bulk soil C increased relative to treeless controls.

Several features of our site likely made early bulk soil C responses difficult to detect. Former pasture soils can have high baseline soil C, and new tree-derived inputs are offset by losses or turnover of legacy C, resulting in little net change in the total bulk soil C pool during early reforestation (Marín-Spiotta et al., 2009; Nave et al., 2013; O’Kelley et al., 2025). Establishment activities also contribute to short-term C losses or variability by disturbing soil structure and exposing protected OM to decomposition, particularly in the first years after planting (Paul et al., 2002). Although survivorship among some species was significantly lower than others, this did not affect our analyses of soil C because we measured soil on an individual tree basis next to healthy saplings rather than in a larger reforestation area. Additionally, because both tree and control samples in our study were collected from within planting furrows, any effects of initial furrow disturbance were part of the shared site context rather than a confounding difference between treatments.

Bulk soil C alone provides limited information about whether soil C is vulnerable to decomposition or positioned within structures that promote persistence (O’Rourke et al., 2015; Schmidt et al., 2011). Aggregate-associated C and aggregate stability added this context in our study by showing that reforestation effects were expressed through physical fractions that differ in accessibility, turnover, and potential protection (Jastrow et al., 2007; Six et al., 2000). Experimental evidence supports this interpretation, as aggregate disruption can increase C mineralization and aggregate turnover can regulate both decomposition of new inputs and transfer of C into smaller, more protected fractions (Kpemoua et al., 2022; Liu et al., 2023). Viewed in this context, the treatment differences in aggregate-associated C and MWD suggest that early reforestation altered soil C organization even though bulk soil C did not increase relative to controls. For restoration monitoring, these metrics can therefore reveal early changes in belowground structure and C distribution that bulk soil C alone may miss.

### Surface macroaggregates captured early structural change

Early reforestation effects in our study were concentrated in surface macroaggregates (>250 µm), the fraction that held most aggregate-associated C and showed clearer treatment differences than bulk soil C. In longer-term vegetation recovery studies, macroaggregates often show strong surface-soil responses, including increases in abundance, stability, and associated C (Hu & Lan, 2020; Shi et al., 2023; Wang, Zhong et al., 2020). This responsiveness reflects both their position in the soil profile and their formation pathways. Surface mineral soil receives concentrated inputs from fine roots and rhizodeposition, which stimulate microbial activity and contribute to soil C formation (Rasse et al., 2005; Sokol et al., 2019). Macroaggregates are stabilized by relatively labile organic inputs, roots, and microbial binding agents, making them closely tied to recent plant and microbial activity (Jastrow et al., 2007; Six et al., 2004; Tisdall & Oades, 1982). Our results suggest that similar surface macroaggregate responses can begin to differentiate soils within the first few years of reforestation.

Across recovery chronosequences, increases in macroaggregate abundance, stability, and macroaggregate-associated C track bulk SOC gains during forest and vegetation recovery (Hu & Lan, 2020; Shi et al., 2023; Wang, Zhong et al., 2020). However, the early macroaggregate C responses observed here should not be interpreted as clear evidence of long-term C stabilization. Three years after reforestation, greater C in larger aggregate fractions indicates redistribution of C within dynamic soil structure rather than persistent C accumulation in stabilized pools. Macroaggregates are important transitional fractions because they receive fresh plant-derived C and undergo relatively rapid formation and turnover, creating opportunities for C to be decomposed, retained, or transferred into smaller aggregate fractions (Oades, 1984; Six et al., 2004). Longer-term stabilization depends on whether C is ultimately incorporated into microaggregates and mineral-associated fractions where microbial access is more restricted (Six et al., 2000; Six et al., 2004). The weak treatment responses in microaggregates and silt-and-clay fractions suggest that reforestation reorganized soil C within recent and structurally dynamic pools, but had not yet produced clear evidence of persistent increases in stabilized soil C.

### Species responses were consistent with mycorrhizal and pasture-legacy mechanisms

The surface macroaggregate response was not uniform across planted species. Macroaggregate proportion, stability, and aggregate-associated C remained closer to control values under ponderosa pine (*Pinus ponderosa*, [EcM]) than under incense cedar (*Calocedrus decurrens* [AM]) or black cottonwood (*Populus trichocarpa* [AM/EcM]), suggesting that early aggregate development reflected species-specific differences in root- and fungi-mediated soil structure rather than a uniform effect of tree planting. The weaker ponderosa pine response was broadly consistent with our expectation that AM species would show stronger early aggregate responses than the EcM pine under former-pasture legacy conditions. Mycorrhizal inoculation studies support the role of fungal symbionts in these aggregate responses, showing that AM fungi increase rhizosphere macroaggregates and aggregate stability (Li et al., 2026; Zhang et al., 2019), while both AM and EcM fungi can increase aggregate stability when compatible host-fungal associations establish successfully (Demenois et al., 2017; Graf and Frei, 2013). Thus, the weaker ponderosa pine response likely reflects limited compatibility with former-pasture soil conditions rather than a lower inherent capacity of EcM fungi to promote aggregation.

Importantly, black cottonwood can associate with both AM and EcM fungi, so its aggregate responses should not be interpreted as evidence of a simple AM-versus-EcM divide. The similarity between black cottonwood and incense cedar in surface aggregate stability and large macroaggregate proportion suggests that early aggregate responses may have reflected compatibility with AM belowground conditions during establishment. This interpretation is consistent with evidence that *Populus* fungal associations change through early development, with EcM fungi increasing over time while AM colonization may persist rather than disappear entirely (Argiroff et al., 2024; Nash et al., 2025).

Former pasture provides the landscape context for this compatibility interpretation because reforestation introduced tree seedlings into soils recently shaped by herbaceous vegetation with AM fungal communities. In natural recovery, shifts from grassland or shrubland toward forest are often accompanied by changes in dominant mycorrhizal strategies, with AM plants common in earlier stages and EcM trees becoming more prominent during later forest development. Along a grassland-to-primary forest succession, mycorrhizal controls on aggregation changed through time, with AM fungal communities especially important during grassland and shrubland stages and different aggregation controls emerging in later EcM forest stages (Qiang et al., 2023). Land-use history can further alter this trajectory. In recovering eastern U.S. forests, agricultural abandonment was associated with a six-fold increase in AM tree dominance relative to historical forest composition, suggesting that former agricultural land use can leave a long-lasting imprint on mycorrhizal composition during forest recovery (Wurzburger et al., 2023). Our findings suggest that former pasture may favor early aggregate development by AM-compatible species while slowing the development of EcM-mediated belowground pathways.

The weaker early aggregate response under ponderosa pine could reflect limited compatibility with inherited soil biotic conditions, including reduced access to compatible EcM inoculum. Although this site was historically forested and located near intact forest across the McKenzie River, approximately 60 years of fallow pasture conditions may have reduced or filtered compatible EcM inoculum. EcM inoculum can persist in soil for years to decades, but persistence is often taxonomically limited and shaped by prior host presence (Glassman et al., 2015; Nguyen et al., 2012; Shemesh et al., 2023). EcM inoculum may also arrive from nearby intact forest through aerial spore dispersal, but dispersal and seedling colonization generally decline with distance from source forests (Galante et al., 2011; Peay et al., 2012). Such limits to compatible EcM inoculum could affect not only aggregation directly, but also early tree establishment, which determines the plant inputs available to drive belowground change.

This connection between mycorrhizal compatibility and tree establishment is consistent with the survivorship pattern across the broader planting. EcM ponderosa pine had lower survivorship than both AM incense cedar and AM/EcM black cottonwood, suggesting that former pasture conditions may have been less favorable for rapid EcM tree establishment. Mycorrhizal compatibility can affect early regeneration outcomes, as AM seedlings have higher survival in AM legacy plots following timber harvest (Fitch et al., 2026), and low EcM inoculum can reduce establishment and growth of introduced Pinaceae, including ponderosa pine, away from existing plantations (Nuñez et al., 2009). Survivorship differences could also reflect species-specific tolerance to planting stress, first-year soil moisture stress, or other establishment constraints. However, compatible mycorrhizal associations can mediate plant responses to stressful conditions by improving water uptake, drought recovery, survival, and growth under stress (Lehto & Zwiazek, 2011; Pereira et al., 2021; Marro et al., 2022). Survivorship is significant for soil C trajectories because dead seedlings contribute less root biomass, rhizodeposition, and litter input than successfully established trees over time. Because root traits, rhizodeposition, and mycorrhizal association help determine how plant inputs enter SOM stabilization pathways, pasture legacy may influence belowground development both directly through tree survival and productivity and indirectly through root- and fungal-mediated soil structure (Hamza et al., 2026; Poirier et al., 2018).

Mycorrhizal compatibility did not fully explain the species-level differences in aggregate and C responses. Incense cedar and black cottonwood both showed stronger surface aggregate responses than ponderosa pine, but they differed from each other in aggregate-associated C and bulk soil C. This divergence was clearest for black cottonwood, which had high surface-soil MWD and large macroaggregate abundance without proportional C accumulation. Both incense cedar and black cottonwood showed positive relationships between MWD and bulk soil C, but only black cottonwood showed a positive relationship between MWD and bulk soil C:N. Because C:N ratios are shaped by litter and root chemistry and decomposition, the black cottonwood pattern suggests that aggregate stability was linked to variation in plant inputs or C turnover within this treatment, rather than to C pool size alone (Manzoni et al., 2010; Lorenz et al., 2020). One possible explanation is that black cottonwood promoted both aggregate formation and aggregate turnover, allowing rapid structural reorganization while also stimulating C loss through decomposition. Root growth and turnover can physically disturb existing aggregates and expose protected C to decomposition (Dijkstra et al., 2021). Rhizosphere priming provides another pathway, as living roots can stimulate SOM decomposition through exudation (Huo et al., 2017; Kuzyakov, 2002), and stronger priming has been linked experimentally to faster aggregate turnover (Wang, Yin, et al., 2020). Black cottonwood is widely recognized as a fast-growing riparian tree (Geraldes et al., 2014) and was the tallest sampled species in our study. As a broadleaf deciduous species, black cottonwood also differs from the two evergreen conifers in the timing of leaf inputs, and litterfall phenology, which can influence decomposition dynamics (Pearse et al., 2014) and microbial decomposition rates (Makita & Fujii, 2015).

These results do not suggest that black cottonwood is unfavorable for soil recovery. Rather, black cottonwood may represent a faster-turnover pathway in which soil structure changes before net C gains become apparent. For monitoring, this distinction is important because MWD reflects aggregate size distribution and structural stability, not net C storage directly. Early increases in aggregate stability should therefore be interpreted alongside aggregate-associated C, bulk soil C, and indicators of C processing.

### δ^13^C patterns supported aggregate hierarchy expectations

In our study, smaller aggregate fractions were progressively enriched in δ¹³C, supporting the aggregate hierarchy framework by indicating that smaller, more protected fractions contained C that was more microbially processed than C in larger, more dynamic fractions. This interpretation is consistent with studies showing that δ¹³C commonly increases with decomposition, soil depth, and increasing microbial contribution to SOM, as microbes preferentially respire ¹²C and microbial residues become increasingly represented in persistent soil fractions (Acton et al., 2013; Klink et al., 2022; Lorenz et al., 2020). Aggregate and fractionation studies show similar patterns, with δ¹³C enrichment increasing from macroaggregates or particulate fractions toward microaggregates, mineral-associated fractions, or other physically protected pools (Gunina & Kuzyakov, 2014; Liu et al., 2018; Scartazza et al., 2023). The pattern is also consistent with a model in which relatively processed microaggregate and mineral-associated materials serve as components of larger, more dynamic aggregates (Silva et al., 2015). The accompanying decline in aggregate-fraction C:N provides complementary stoichiometric evidence for this fraction-level processing gradient (Lorenz et al., 2020; Scartazza et al., 2023). Together, these patterns provide mechanistic support that aggregate fractions differ in C processing and protection.

The lack of treatment-level δ¹³C and aggregate-fraction C:N differences suggests that species effects on C processing and stabilization had not yet clearly emerged after three years, even though species differences were already evident in aggregate structure and C distribution. Natural-abundance δ¹³C reflects both plant source signatures and decomposition history, so treatment-level differences are most likely to emerge when species inputs have distinct isotope signatures or when enough time has passed for differences in microbial processing to accumulate (Krüger et al., 2024; Lorenz et al., 2020). Longer-term sampling, paired with direct measurements of litter, roots, and mycorrhizal isotope signatures, would be needed to determine whether species-level isotope differences emerge as reforestation proceeds. Thus, the isotope results support the interpretation that early reforestation effects were expressed first through structural reorganization and aggregate C distribution, while species-level differences in stabilized C may require more time to develop.

### Applications for restoration monitoring

Our results suggest that aggregate-based measures can help identify belowground divergence among plantings before bulk soil C stocks provide a clear signal. Restoration monitoring frameworks emphasize that indicators should be tied to project goals, sensitive to expected recovery processes, and useful for evaluating recovery trajectories over time (Gann et al., 2019; Prach et al., 2019). Aggregate stability, aggregate size distribution, and aggregate-associated C meet this need by showing whether plantings are altering soil structure, C distribution, and potential C protection pathways without assuming that these changes already represent net soil C sequestration. This distinction is especially important in young reforestation projects, where monitoring windows are often shorter than the time required to detect bulk soil C gains.

Aggregate-based measures also broaden monitoring beyond C storage alone. Soil aggregation influences erosion resistance and infiltration, so aggregate stability is relevant where restoration goals include soil protection, hydrologic recovery, or resilience to more variable precipitation (Barthès & Roose, 2002; Bronick & Lal, 2005). These functions will become increasingly important in reforestation projects facing drought stress, variable precipitation, or more intense rainfall under climate change. Because aggregation is biologically mediated, aggregate responses can also indicate whether plantings are developing root- and fungal-mediated soil structure (Rillig & Mummey, 2006; Wilpiszeski et al., 2019). In practice, aggregate stability and size distribution can serve as screening metrics because they require relatively low-cost wet-sieving equipment, whereas aggregate-associated C and δ¹³C provide more detailed but more analytically intensive information about C distribution and processing.

Beyond individual sites, standardized aggregate measurements could help restoration programs compare early belowground responses across species, soil conditions, and land-use histories. Early aggregate responses can help identify which tree species and site conditions promote belowground change, allowing restoration programs to refine planting designs before long-term soil C outcomes are measurable. This kind of indicator-based learning is already used in agricultural soil-health assessment, where aggregate stability helps track management effects across broad site networks and identify soil structural responses not captured by other common soil-health measures (Bagnall et al., 2023; Blair et al., 2024). In restoration settings, a similar approach could help determine whether species-specific belowground responses are consistent across sites or depend on land-use legacies such as former pasture conditions and inherited mycorrhizal communities.

This cross-site perspective is especially relevant in regions where reforestation is expanding across former agricultural or pasture lands, because inherited soil conditions may influence both tree establishment and the early aggregate-mediated pathways through which plant inputs enter soil C pools (Hamza et al., 2026; Nuñez et al., 2009). In the Pacific Northwest, where riparian reforestation is increasingly used to restore ecosystem function and increase carbon storage, aggregate-based monitoring could help place individual projects within a broader regional understanding of soil recovery trajectories (O’Kelley et al., 2025). Future work should follow these plots through time and pair aggregate measurements with direct assessments of plant inputs, mycorrhizal development, and soil C stocks to test whether early aggregate shifts predict later increases in stabilized C and bulk soil C storage.

Overall, this study shows that three years after reforestation on former pasture, belowground change was detectable in aggregate stability, aggregate size distribution, and aggregate-associated C even though bulk soil C had not increased relative to treeless controls. Aggregate-based measures therefore provide practical early indicators of reforestation trajectories and can help clarify how tree species identity and land-use legacy shape soil C organization in young plantings.

## Acknowledgements

We acknowledge the Eugene Water and Electric Board for their funding of this work and of the experimental reforestation site. Additional funding came from the Environmental Science Summer Research Fellowship of the Environmental Studies Program at the University of Oregon. We thank Matt Fehrenbacher of Trout Mountain Forestry for leading the reforestation efforts and maintaining the site with the help of Franco Reforestation Inc. We express our gratitude to the following individuals for their assistance with field and laboratory work: Sarah Weber, Tetianna Smith-Drysdale, Ethan Torres, and the 2023 undergraduate student cohort of the Environmental Leadership Program’s Natural Climate Solutions team at the University of Oregon.

## Author Contributions

CEM conceptualized and designed the soil aggregate experiment with input and supervision from LCRS. HRD and ERH conceptualized and designed the experimental reforestation with input and supervision from LCRS, performed fieldwork, and provided support with experimental design of the study. CEM and ERH performed fieldwork for sample collection. CEM conducted laboratory and statistical analyses, and drafted the original manuscript. This study was funded by the Eugene Water and Electric Board (acquired by LCRS) and through the University of Oregon’s Environmental Science Summer Research Fellowship (acquired by CEM). All authors reviewed and edited the final manuscript.

