## Appendix S1- Tables and Figures for "Soil aggregates reveal tree species and land-use legacy effects on early carbon storage pathways during reforestation"

**Table S1:** Mean aggregate-associated organic carbon (g C $kg^{-1}$ bulk soil) by treatment and soil depth, with one-way ANOVA results for treatment differences within each depth × fraction combination.

| Depth (cm) | Aggregate fraction | Control | Incense cedar | Black cottonwood | Poderosa pine | *F* value | *p* value |
|---|---|---|---|---|---|---|---|
| 0-20 | macro | 6.88 ± 1.13 | 11.3 ± 1.3 | 10.79 ± 1.56 | 6.37 ± 1 | 4.088 | **0.0109*** |
| 0-20 | meso | 11.02 ± 0.77 | 10.93 ± 0.9 | 7.06 ± 0.77 | 11.44 ± 0.74 | 6.69 | **0.0006***** |
| 0-20 | micro | 3.84 ± 0.26 | 3.33 ± 0.33 | 2.22 ± 0.21 | 4 ± 0.34 | 7.687 | **0.0002***** |
| 0-20 | silt & clay | 0.77 ± 0.07 | 0.72 ± 0.1 | 0.42 ± 0.06 | 0.82 ± 0.08 | 5.335 | **0.0027**** |
| 20-40 | macro | 4.45 ± 0.84 | 4.07 ± 1.25 | 3.83 ± 1.16 | 2.42 ± 0.52 | 0.785 | 0.5070 |
| 20-40 | meso | 9.32 ± 0.87 | 10.39 ± 0.83 | 8.67 ± 0.44 | 11.46 ± 1.01 | 2.318 | 0.0858 |
| 20-40 | micro | 3.63 ± 0.32 | 4.13 ± 0.48 | 2.62 ± 0.3 | 3.84 ± 0.4 | 2.979 | **0.0394*** |
| 20-40 | silt & clay | 0.68 ± 0.08 | 0.84 ± 0.09 | 0.47 ± 0.04 | 0.79 ± 0.04 | 5.852 | **0.0016**** |

**Table S1 notes:** Values are means ± standard error of aggregate-fraction carbon content (g C $kg^{-1}$ bulk soil). *F* and *p* values are from one-way ANOVAs testing for treatment differences within each depth × fraction combination. Boldface *p* values indicate significant treatment effects ($p < 0.05$).

**Table S2:** Mean fraction-specific organic carbon concentration (%C) by treatment and soil depth, with one-way ANOVA results for treatment differences within each depth × fraction combination.

| Depth (cm) | Aggregate fraction | Control | Incense cedar | Black cottonwood | Poderosa pine | *F* value | *p* value |
|---|---|---|---|---|---|---|---|
| 0-20 | macro | 2.60 ± 0.13 | 3.02 ± 0.09 | 2.36 ± 0.06 | 2.74 ± 0.14 | 6.312 | **0.0009*** ** |
| 0-20 | meso | 2.42 ± 0.15 | 2.65 ± 0.1 | 2.18 ± 0.14 | 2.35 ± 0.13 | 2.335 | 0.0840 |
| 0-20 | micro | 2.08 ± 0.15 | 2.43 ± 0.09 | 1.67 ± 0.08 | 2.19 ± 0.1 | 9.201 | **<0.0001*** ** |
| 0-20 | silt & clay | 2.26 ± 0.11 | 2.60 ± 0.11 | 1.99 ± 0.06 | 2.44 ± 0.09 | 8.311 | **0.0001*** ** |
| 20-40 | macro | 2.37 ± 0.12 | 2.59 ± 0.12 | 2.13 ± 0.1 | 2.61 ± 0.21 | 2.484 | 0.0705 |
| 20-40 | meso | 1.9 ± 0.14 | 2 ± 0.15 | 1.6 ± 0.1 | 1.95 ± 0.12 | 2.019 | 0.1220 |
| 20-40 | micro | 1.79 ± 0.16 | 1.99 ± 0.13 | 1.34 ± 0.06 | 1.88 ± 0.09 | 6.687 | **0.0006*** ** |
| 20-40 | silt & clay | 2.04 ± 0.15 | 2.29 ± 0.12 | 1.75 ± 0.07 | 2.2 ± 0.07 | 5.229 | **0.0031** ** |

**Table S2 notes:** Values are means ± standard error of aggregate-fraction carbon concentration (%C). *F* and *p* values are from one-way ANOVAs testing for treatment differences within each depth × fraction combination. Boldface *p* values indicate significant treatment effects ($p < 0.05$).

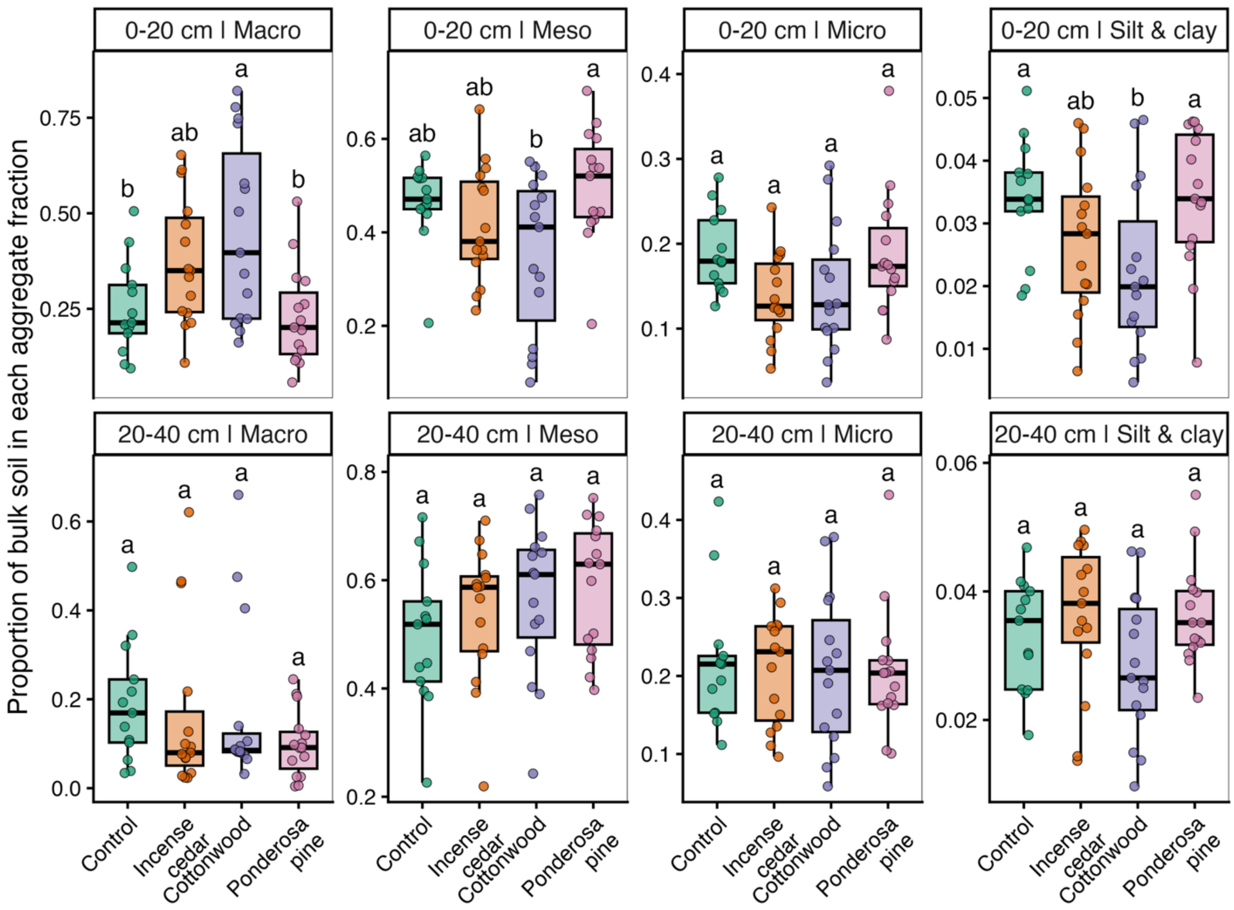


**Figure S1:** Aggregate-size proportions by treatment and depth. Boxplots show the mass proportion of bulk soil in each aggregate fraction for macroaggregate, mesoaggregate, microaggregate, and silt-and-clay fractions in the 0-20 cm and 20-40 cm soil layers. Letters indicate significant treatment differences within each depth × fraction combination based on pairwise Tukey's HSD tests performed on logit-transformed proportion values, while raw proportion values are shown for interpretability.

**Table S3:** Mean aggregate-fraction C:N by treatment and soil depth, with one-way ANOVA results for treatment differences within each depth × fraction combination.

| Depth (cm) | Aggregate fraction | Control | Incense cedar | Black cottonwood | Ponderosa pine | *F* value | *p* value |
|---|---|---|---|---|---|---|---|
| 0-20 | macro | 11.24 ± 0.28 | 11.42 ± 0.24 | 11.19 ± 0.22 | 10.4 ± 0.5 | 1.911 | 0.1388 |
| 0-20 | meso | 11.21 ± 0.24 | 11.35 ± 0.22 | 11.11 ± 0.28 | 10.92 ± 0.24 | 0.564 | 0.6414 |
| 0-20 | micro | 10.71 ± 0.23 | 11.03 ± 0.24 | 10.53 ± 0.31 | 10.89 ± 0.17 | 0.796 | 0.5014 |
| 0-20 | silt & clay | 10.59 ± 0.22 | 10.98 ± 0.47 | 9.99 ± 0.33 | 10.64 ± 0.2 | 1.449 | 0.2394 |
| 20-40 | macro | 11.35 ± 0.27 | 11.77 ± 0.35 | 10.76 ± 0.12 | 11.42 ± 0.35 | 2.169 | 0.1023 |
| 20-40 | meso | 11.3 ± 0.21 | 11.41 ± 0.41 | 11.21 ± 0.26 | 11.13 ± 0.27 | 0.16 | 0.9225 |
| 20-40 | micro | 11.19 ± 0.41 | 11.05 ± 0.3 | 10.5 ± 0.37 | 10.75 ± 0.2 | 0.897 | 0.4486 |
| 20-40 | silt & clay | 11.46 ± 0.45 | 11.47 ± 0.41 | 10.33 ± 0.26 | 10.64 ± 0.16 | 2.99 | **0.0389*** |

**Table S3 notes:** Values are means ± standard error of aggregate-fraction C:N. $F$ and $p$ values are from one-way ANOVAs testing for treatment differences within each depth × fraction combination. Boldface $p$ values indicate significant treatment effects ($p < 0.05$).

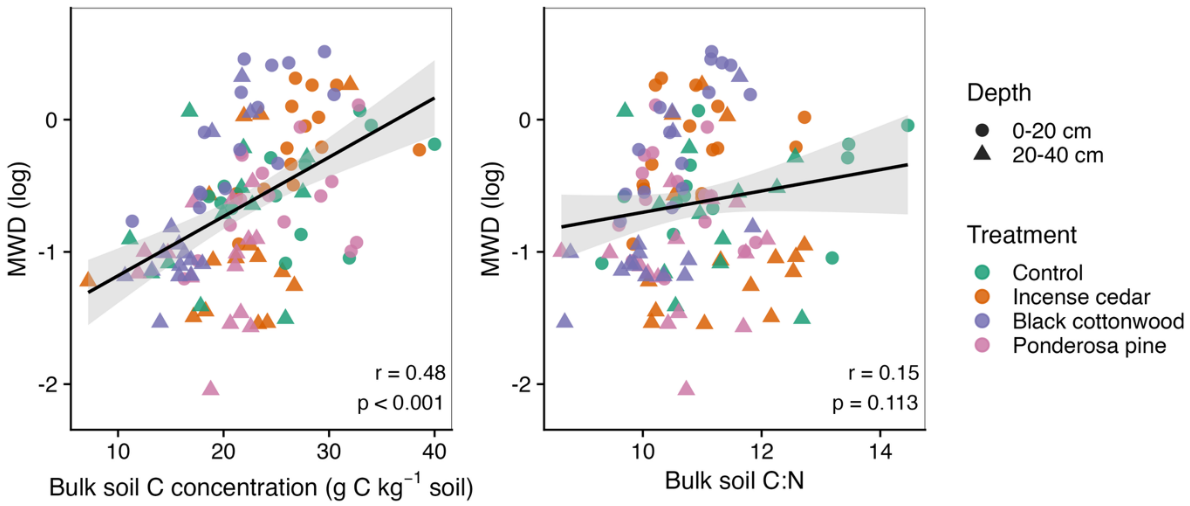


**Figure S2:** Relationships between log-transformed mean weight diameter (MWD) and bulk soil C concentration or bulk soil C:N across the full dataset. Points are colored by treatment and shaped by soil depth; black lines show overall linear fits with 95% confidence intervals. Pearson correlation coefficients (*r*) and associated *p* values are shown within each panel. Because the strength of both relationships differed significantly among treatments, treatment-specific slopes are shown in Figure 2B of the main text.

**Table S4:** Mean bulk soil C concentration, bulk soil N concentration, and bulk soil C:N by treatment and soil depth, with one-way ANOVA results for treatment differences within each depth × measure combination.

| Depth (cm) | Bulk soil measure | Control | Incense cedar | Black cottonwood | Ponderosa pine | *F* value | *p* value |
|---|---|---|---|---|---|---|---|
| 0-20 | C | 26.8 ± 1.79 | 26.92 ± 1.13 | 21.83 ± 1.29 | 25.16 ± 1.39 | 2.928 | **0.0418*** |
| 0-20 | N | 2.34 ± 0.12 | 2.48 ± 0.1 | 2.03 ± 0.1 | 2.37 ± 0.11 | 3.314 | **0.0266*** |
| 0-20 | C:N | 11.46 ± 0.45 | 10.88 ± 0.22 | 10.66 ± 0.17 | 10.58 ± 0.18 | 2.099 | 0.1111 |
| 20-40 | C | 20.1 ± 1.59 | 21.63 ± 1.42 | 16.4 ± 0.79 | 19.22 ± 0.97 | 3.507 | **0.0214*** |
| 20-40 | N | 1.79 ± 0.12 | 1.9 ± 0.13 | 1.61 ± 0.07 | 1.8 ± 0.08 | 1.603 | 0.1996 |
| 20-40 | C:N | 11.2 ± 0.27 | 11.35 ± 0.24 | 10.2 ± 0.23 | 10.63 ± 0.24 | 4.863 | **0.0046*** |

**Table S4 notes:** Values are means ± standard error. Bulk soil C and N are reported as g $kg^{-1}$ soil. *F* and *p* values are from one-way ANOVAs testing for treatment differences within each depth × bulk soil measure combination. Boldface *p* values indicate significant treatment effects ($p < 0.05$).

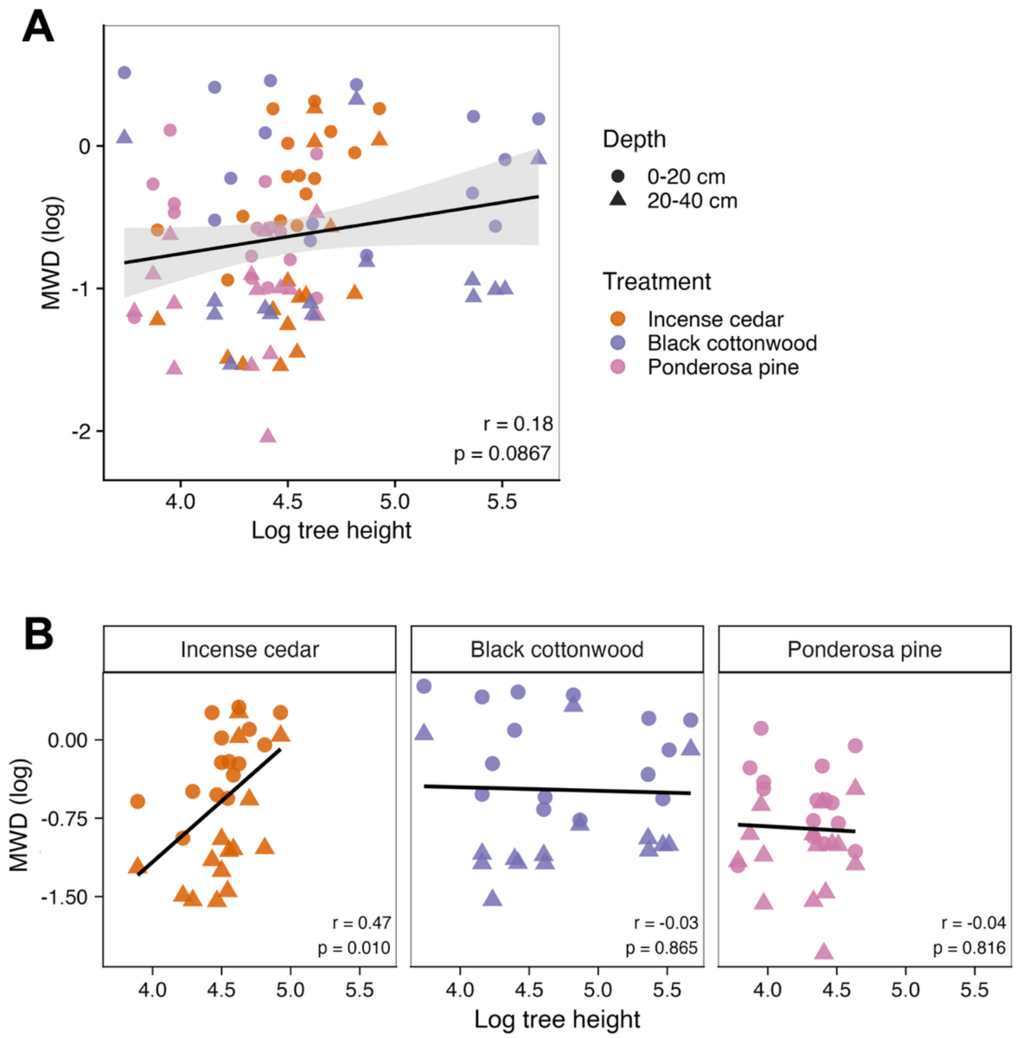


**Figure S3:** Relationship between log-transformed mean weight diameter (MWD) and log-transformed tree height for sampled planted trees. Control samples are not included because tree height was measured only for planted individuals. (A) Overall relationship across incense cedar, black cottonwood, and ponderosa pine soils. (B) Species-specific relationships. Points are colored by tree species and shaped by soil depth; black lines show linear fits with 95% confidence intervals in panel A and species-specific linear fits in panel B. Pearson correlation coefficients (*r*) and associated *p* values are shown within each panel.
